# Single-cell RNA sequencing of a European and an African lymphoblastoid cell line

**DOI:** 10.1101/548115

**Authors:** Daniel Osorio, Xue Yu, Peng Yu, Erchin Serpedin, James J. Cai

**Affiliations:** Department of Veterinary Integrative Biosciences, Texas A&M University, College Station, TX 77843, USA; Department of Electrical and Computer Engineering, Texas A&M University, College Station, TX 77843, USA; Interdisciplinary Program of Genetics, Texas A&M University, College Station, TX 77843, USA

## Abstract

In biomedical research, lymphoblastoid cell lines (LCLs), often established by *in vitro* infection of resting B cells with Epstein Barr Virus, are commonly used as surrogates for peripheral blood lymphocytes. Genomic and transcriptomic information on LCLs has been used to study the impact of genetic variation on gene expression in humans. Here we present single-cell RNA sequencing (scRNA-seq) data on GM12878 and GM18502—two LCLs derived from the blood of female donors of European and African ancestry, respectively. Cells from three samples (the two LCLs and a 1:1 mixture of the two) were prepared separately using a 10X Genomics Chromium Controller and deeply sequenced. The final dataset contained 7,045 cells from GM12878, 5,189 from GM18502, and 5,820 from the mixture, offering valuable information on single-cell gene expression in highly homogenous cell populations. This dataset is a suitable reference of population differentiation in gene expression at the single-cell level. Data from the mixture provides additional valuable information facilitating the development of statistical methods for data normalization and batch effect correction.

## Background & Summary

Immortalized cell lines are continuously growing cells derived from biological samples. As long-lasting supplies of cells containing genotypic and phenotypic information matching that of their origins, cell lines have contributed significantly to biomedical research. Lymphoblastoid cell lines (LCLs) are one of important members among many immortalized cell lines^1^. LCLs are usually established by infecting human peripheral blood lymphocytes in vitro with Epstein Barr Virus (EBV). The infection selectively immortalizes resting B cells, giving rise to an actively proliferating B cell population^2^. LCLs exhibit a low somatic mutation rate in continuous culture, making them the preferred choice of storage for individual’s genetic material^3^. As one of the most reliable, inexpensive, and convenient sources of cells, LCLs have been used by several large-scale genomic DNA sequencing efforts such as the International HapMap and the 1,000 Genomes projects, in which a large collection of LCLs were derived from individuals of different genetic backgrounds, to document the extensive genetic variation in human populations^4,5^.

LCLs are also an in vitro model system for a variety of molecular and functional assays, contributing to studies in immunology, cellular biology, genetics, and many other research areas^6-12^. It is also believed that gene expression in LCLs encompasses a wide range of metabolic pathways specific to individuals where the cells originated^13^. LCLs have been used in population-scale RNA sequencing projects^14-16^, as well as epigenomic projects^17^. For many reference LCLs, both genomic and transcriptomic information is available, making it possible to detect the correlation between genotype and expression level of genes and infer the potential causative function of genetic variants^18^. Furthermore, comparisons of gene expression profiles of LCLs between populations such as between Centre d’Etude du Polymorphisme Humain – Utah (CEPH/CEU) and Yoruba in Ibadan, Nigeria (YRI), have revealed the genetic basis underlying the differences in transcriptional activity between the two populations^16,19^.

With the advent of single-cell RNA sequencing (scRNA-seq) technology^20,21^, our approach for understanding the origin, global distribution, and functional consequences of gene expression variation needs to be upgraded. Data generated from scRNA-seq provides an unprecedented resolution of the gene expression profiles at single cell level, which allows the identification of previously unknown subpopulations of cells and functional heterogeneity in a cell population^22-24^.

In this study, we used scRNA-seq to measure gene expression across thousands of cells from two LCLs: GM12878 and GM18502. Cells were prepared using a 10X Genomics Chromium Controller (10X Genomics, Pleasanton, CA) as described previously^21^ and sequenced using an Illumina Novaseq 6000 sequencer. We present this dataset on the single-cell gene expression profile of 7,045 cells from GM12878 and 5,189 from GM18502. GM12878 is a popular sample that has been widely used in genomic studies. For example, it is one of three ‘Tier 1’ cell lines of the Encyclopedia of DNA Elements (ENCODE) project^17,25^. GM18502, derived from the donor of African ancestry, serves as a representative sample from the divergent population. The two cell lines are part of the International HapMap project, and genotypic information is available for both of them^4^. We also processed and sequenced an additional sample of 1:1 mixture of GM12878 and GM18502 using the same scRNA-seq procedure. Our dataset presented here provides a suitable reference for those researchers interested in performing between-populations comparisons in gene expression at the single-cell level, as well as for those developing new statistical methods and algorithms for scRNA-seq data analysis.

## Methods

### Cell Culture

GM12878 and GM18502 were purchased from the Coriell Institute for Medical Research. Cells were cultured in the Roswell Park Memorial Institute (RPMI) Medium 1640 supplemented with 2mM L-glutamine and 20% of non-inactivated fetal bovine serum in T25 tissue culture flasks. Flasks with 20 mL medium were incubated on the upright position at 37°C under 5% of carbon dioxide. Cell cultures were split every three days for maintenance.

### Growth curve

Four culture flasks for each cell line were started with 200,000 viable cells/mL to measure the growth rate of each cell line. Cells were prepared and cultured as described above. Viable cell number was estimated on a daily basis for four days using Trypan Blue staining and counted in Neubauer counting chamber (Figure 1a). Briefly, 100 uL suspended cells from each flask were taken every day, to visualize the viable cells, the samples were stained using 10 uL of Trypan Blue (0.4%), and live cells were counted manually using a Neubauer counting chamber.

**Figure 1:**
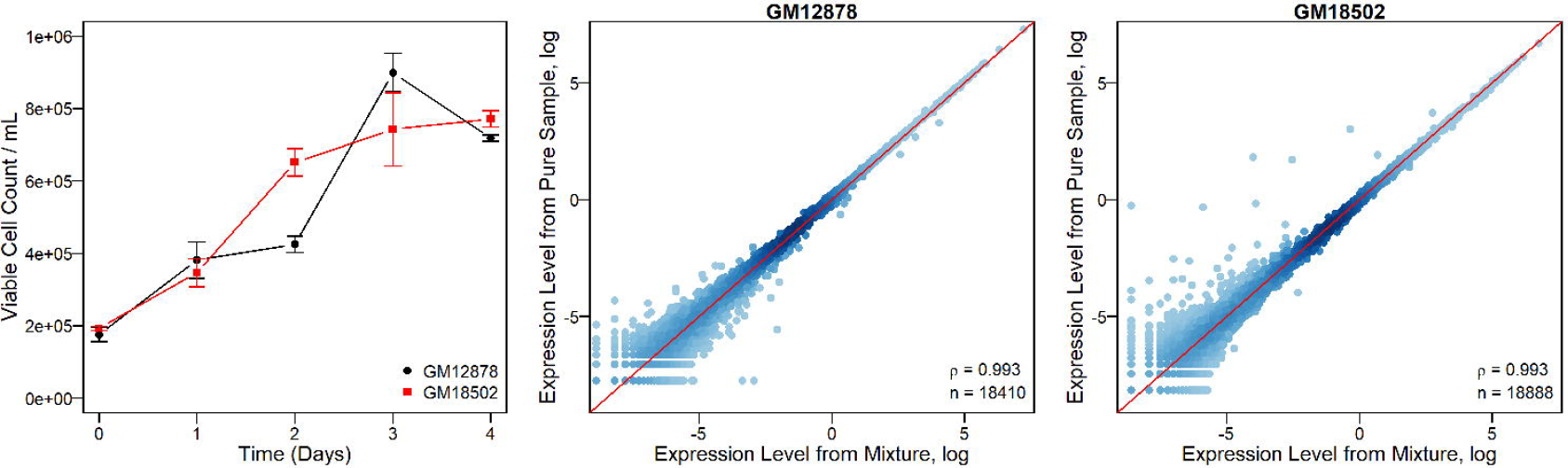
**a.** Growth curve of the GM12878 and GM18502 cultured in the same RPMI 1640 medium. **b.** Correlation between the average expression level of genes in cells of pure CEU (or YRI) cell line and that in CEU (or YRI) cells in the mixture. Values of gene expression are log-transformed, and each dot represents a gene. **c.** Spearman’s correlation between the gene expressions profiles UMI average of the cells assigned to the YRI population from the mixture and those from the GM18502 pure cell line. Values were log-transformed, and each dot represents a gene.

### Single cell preparation

Single-cell sample preparation was conducted according to Sample Preparation Demonstrated Protocol provided by 10X Genomics as follows: 1 mL of cell suspensions from each cell line (day 4, stable phase) was pelleted in Eppendorf tubes by centrifugation (400 g, 5 min). The supernatant was discarded, and the cells pellet was then resuspended in 1X PBS with 0.04% BSA, followed by two washing procedures by centrifugation (150 g, 3 min). After the second wash, cells were resuspended in ~500 uL 1X PBS with 0.04% BSA followed by gently pipetting mix 10-15 times. Cells were counted using an Invitrogen Countess automated cell counter (Thermo Fisher Scientific, Carlsbad, CA) and the viability of cells was assessed by Trypan Blue staining (0.4%).

### Generation of single cell GEMs (Gel bead in EMulsion) and sequencing libraries

Libraries were prepared using the Chromium Controller (10X Genomics, CA) in conjunction with the single-cell 3’ v2 kit. Briefly, the cell suspensions were diluted in nuclease-free water according to manufacturer instructions to achieve a targeted cell count of 5,000 for each cell line. The cDNA synthesis, barcoding and library preparation were then carried out according to the manufacturer’s instructions. Libraries were sequenced in the North Texas Genome Center facilities on a Novaseq 6000 sequencer (Illumina, San Diego). Two output fastq read files for each library were generated; read 1 consist in a 26 bp length read including the cell barcode and a unique molecule identifier (UMI); read 2 is a 98 bp length read that includes the RNA read.

### Mapping of reads to transcripts and cells

Raw reads were demultiplexed using CellRanger (v2.0.0, 10X Genomics) using the ‘`mkfastq`’ in conjunction with ‘`bcl2fastq`’ (v2.17.1.14, Illumina) functions. Reads were aligned to the human reference genome (GRCh38), filtered, and counted using the CellRanger ‘`count`’ command.

### Quality Control

Resulted expression matrix for each cell line and the mixture were loaded and processed using functions in Seurat (v2.3.4) R package^26^ as follows: Expression matrix was loaded using the ‘`Read10X`’ function, the default log-normalization was performed using the ‘`NormalizeData`’ function, followed by a centering and scaling of the normalized values using the ‘`ScaleData`’ function. Common quality control measures for scRNA-Seq including UMI count per cell and percentage of mitochondrial transcripts were calculated (Figure 2) independently for each dataset. The corresponding code used is available online (see Code availability).

**Figure 2:**
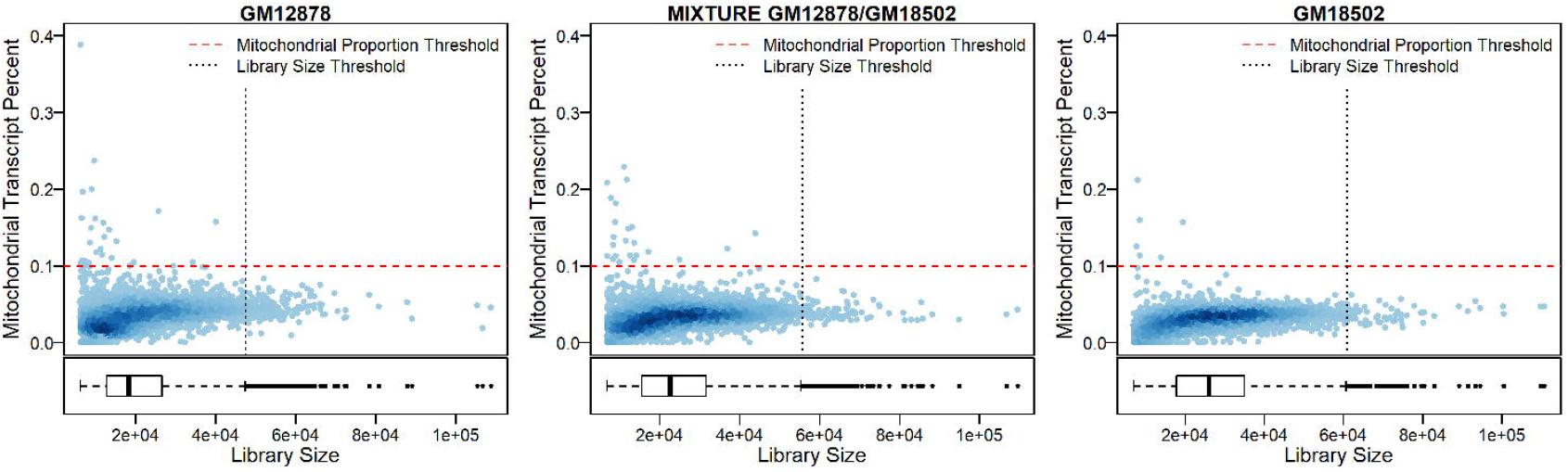
Distribution of the single-cell gene expression profiles under the defined quality control thresholds. There are 6,848 cells for the GM12878, 5,117 for the GM18502 and 5,710 for the mixture sample under the established thresholds.

### Cell cycle phase and population assignment

Cell cycle phase assignment was made using the ‘`CellCycleScoring`’ function included in the Seurat R package, using the marker genes reported for each cell phase in the ‘`cc.genes`’ dataset^27^. Cell population assignment, i.e., assigning cells in the mixture sample back to the cell line (GM12878 or GM18502) they belong to, was made using the Brunet algorithm^28^ for non-negative matrix factorization, included in the NMF (v0.21) R package^29^. A set of marker genes (n = 252) with absolute log-fold change > 2.5 identified by comparing the pure cell lines was chosen as input and the resulting probabilities after 2,000 iterations were used to assign each cell in the mixture to one population. Cells with probabilities equal to 0.5 for each population were reported as 0.

### t-SNE

The set of the more variable genes identified by the function ‘`FindVariableGenes`’ included in the Seurat R package was used as input to produce the t-SNE projection using the ‘`RunTSNE`’ function standard settings (perplexity=30, theta=0.5, maximum iteration=1000, learning rate=250, and momentum reduction=0.5, by using the first 5 components from the principal component analysis).

### Differential expression analysis

Differential expression (DE) analysis was performed using the ‘`FindMarkers`’ function included in the Seurat R package over the cells assigned to the G1 cell cycle phase after normalization. The default Wilcoxon rank sum test^30^ was used, and the minimum log-fold difference was set as 0.25. The associated p-values were corrected using the Bonferroni correction^31^. Functional enrichment analysis of the differentially expressed genes was performed using DAVID, and the identified clusters were filtered by FDR^32^.

### Code availability

All the required code to replicate the characterization of the two cell lines and the mixture, as well as the figures included in this document is available in the sciData-LCL repository (https://github.com/cailab-tamu/sciData-LCL).

## Data Records

The sequencing data from this study has been submitted as the BioProject reference (PRJNA508890), with descriptions of the Biosamples (SUB4895416, SUB4895422, SUB4895423). Raw data of three samples has been deposited at the National Center for Biotechnology Information (NCBI) Sequence Reads Archive (SRA) with accessions: SRS4247443, SRS4238857, and SRS4114895. For each sample, data includes unprocessed scRNA-seq reads in two raw fastq files (*R1.fastq.gz for cell barcodes and UMIs, and *R2.fastq.gz for RNA reads), as well as an expression matrix file in matrix market exchange format (*.mtx) with columns corresponding to cells and row to genes. UMI matrices of this study have been deposited with the Gene Expression Omnibus at GEO: GSE126321. The identifiers for the columns and rows are included in separated files (barcodes.tsv and genes.tsv). These processed files correspond to the output produced by the cell ranger pipeline. In addition, a supplementary table with the barcodes, population, UMI count, gene count, and mitochondrial transcript levels is included.

## Technical Validation

Here we present the gene expression profile for thousands of cells from GM12878 (n=7,045), GM18502 (n=5,189) and a mixture sample of the two (n=5,820). This mixture sample is used as a replicate for both GM12878 and GM18502. The mixture sample also facilitates the direct comparison of gene expression between GM12878 and GM18502 because cells from two cell lines in the mixture were processed simultaneously in the same reaction, eliminating the batch effect. We found that cells in the mixture were assigned back to their cell line origins almost unambiguously using either maker genes (i.e., those expressed in GM12878 but not in GM18502, or vice versa) or non-negative matrix factorization algorithm (see Methods). Furthermore, the average gene expression measured in cells in the mixture, after discriminating cells in the mixture and assigning them to their respective one of original cell lines, was virtually indistinguishable from that measured in the original ‘pure’ cells (Figures 1b and 1c).

The percentage of mitochondrial transcripts, an indicator of apoptotic cell, was computed for all cells sequenced in all the three samples. We found that no more than 0.4% of cells, that is, 26 cells from GM12878, 6 from GM18502, and 23 cells from the mixture sample, surpass the commonly used threshold of 10% mitochondrial transcripts^33^. This suggests that the majority of cells processed and sequenced were viable. Furthermore, as the 10X Genomics Chromium technology relies on droplets to partitioning cells and barcoding, it is normal some of them contain multiple cells in the cell droplet, making the estimation of the frequency of multiplets a critical aspect of quality control^34^. There are several ways to identify multiplets^35-37^. Here we used the 2.5x standard deviation (SD) away from the average library size for each cell as a threshold. Using this threshold, only 171 cells were considered to be multiplets for GM12878, 66 for GM18502, and 87 for the mixture (Figure 2). These results support the quality of the dataset.

In the t-SNE projection, no separation was observed between cells of ‘pure’ lines: GM12878 (Figure 3a) and GM18502 (Figure 3c), and their corresponding replicates in the mixture (Figure 3b). This result suggests that cells in the mixture have the global expression profiles indistinguishable from those of cells of their original samples. Population signal of each sample allows a sample to be separated from others in the first two t-SNE dimensional spaces (Figure 3). Furthermore, for each cell line and the mixture, a continuous path between the different clusters of cells in different cell cycles was identified; this will allow researchers interested in the mechanisms of cell cycle development to perform pseudo-time analysis^38^. Furthermore, for all samples, different clusters of cells in different cell cycles are not completely separated but are linked by a spectrum of cells in intermediate stages; indicating that cell proliferation is a continuous process and researchers interested in this process can use this dataset to refine reference cell populations and calibrate the checkpoints of expression of cell cycle-related genes.

**Figure 3:**
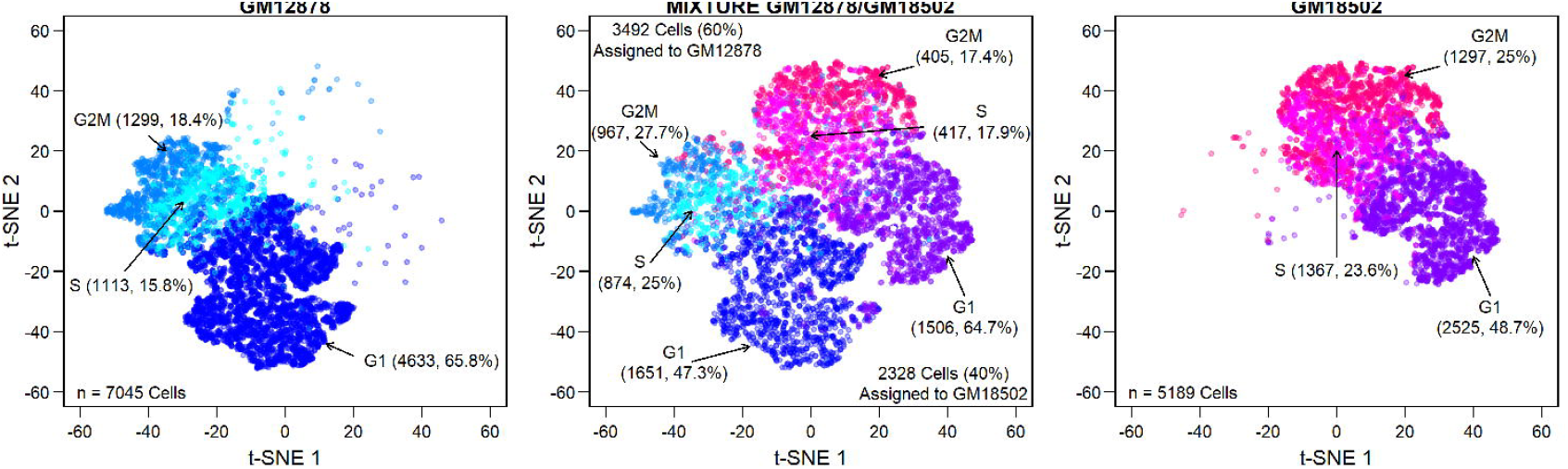
Separate views of t-SNE projection generated from the pooled scRNA-seq data of GM12878 (left panel), GM18502 (right panel), and the mixture (middle panel) samples. Populations and cell cycle phase are labeled and colored differentially.

From the differential gene expression analysis, 153 genes were found overexpressed in the GM12878 cell line and 87 overexpressed in GM18502. Functions of these differentially expressed (DE) genes can be clustered into five significant clusters (FDR < 0.01, corrected from hypergeometric tests) based on the results of DAVID^32^. These significant functional clusters *include immune response, positive regulation of B cell activation, immunoglobulin and major histocompatibility complex composition, cellular chemotaxis, and glycolysis and gluconeogenesis pathways*. Most striking expression differences are observed in immunoglobulin genes such as IGHG3, IGHA1, IGLC3, IGLV3-10 and so on (Figure 4). This highlights the inherited diversity of immune system between two donor individuals. During the formation of the naïve-B cells, gene rearrangement process occurs to reshuffle different subunits of the variable (V), diversity (D) and joining (J) segments of immunoglobulin genes, resulting in the generation of a wide range of organism-specific antigen receptors that allow the immune system to recognize foreign molecules and initiate differential immune responses^39,40^. LCLs are produced through rapid proliferation of few EBV-driven B cells from the blood cell population^41^. Thus, our scRNA-seq data of GM12878 and GM18502 offers a ‘snapshot’ of highly diverse immunoglobulin rearrangement profiles in a much larger population of polyclonal B cells found in the two donors.

**Figure 4:**
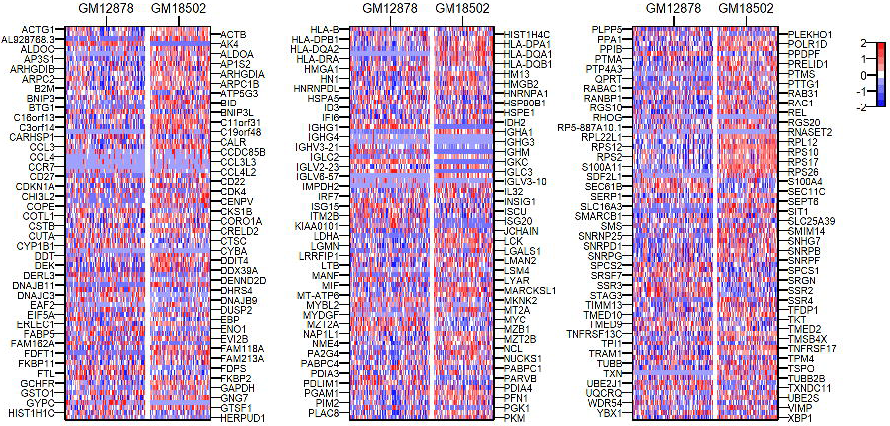
Heatmap of 240 genes highly differentially expressed between GM12878 and GM18502. Data is log-transformed and truncated at a range between −2 and 2 for colouring purposes. Genes are arranged in alphabetical order.

In summary, we present a high-quality dataset of scRNA-seq from homogenous cell populations of two LCLs, including one of the most popular reference cell lines—GM12878. This dataset provides information that can be used to measure cell-to-cell variability in gene expression, study cell status and gene expression changes associated with cell cycles, as well as detect between-population differences in gene expression at the single-cell level. Also, data from the mixture sample is a suitable resource for estimating the technical variability of scRNA-seq and can also be used to calibrate statistical methods for data normalization and batch effect correction.

## Acknowledgements

We thank Andrew Hillhouse for help with single cell preparation and thank Jianhua Huang, Yan Zhong and Guanxiong Li for helpful discussion on data analysis.

## Author contributions

DO, XY, PY, ES and JJC conceived and designed the project; DO and XY cultured the cells; DO and JJC performed bioinformatics analysis, DO, XY, PY, ES and JJC analyzed the data; DO and JJC wrote the manuscript. All authors reviewed the manuscript.

## Manuscript metadata tables

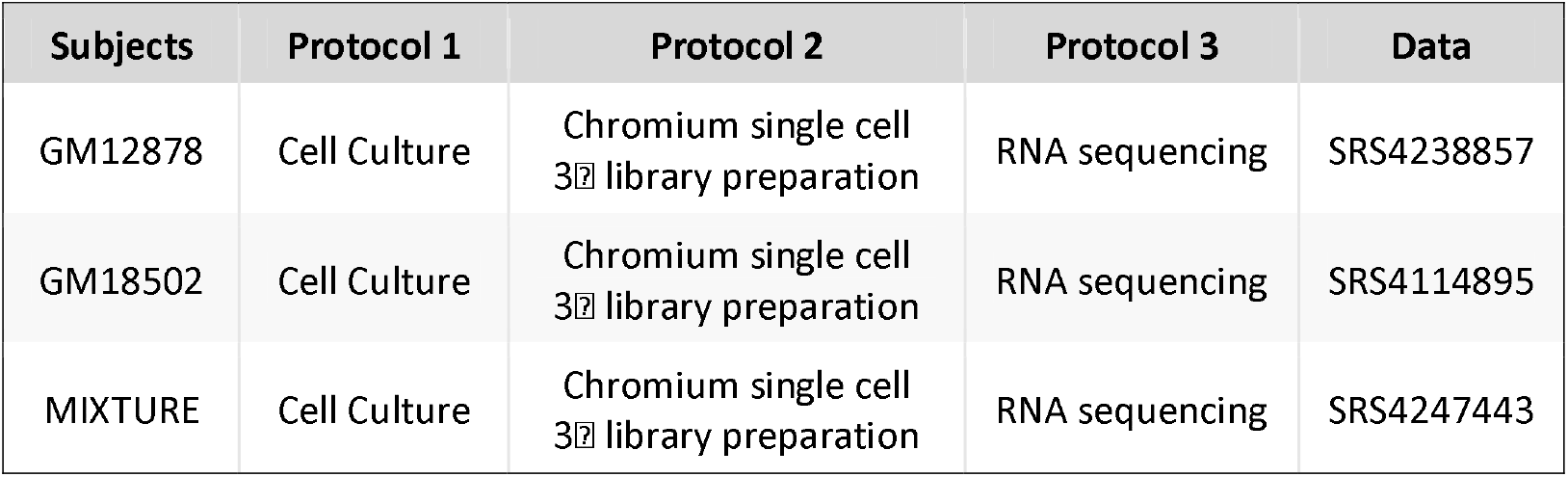

